# Spontaneous Brain Dynamics Associated With Acceleration Of Long-term Functional Connectome In Postnatal Development

**DOI:** 10.1101/2024.11.14.623615

**Authors:** Liang Ma, Sarah Shultz, Zening Fu, Masoud Seraji, Armin Iraji, Vince Calhoun

## Abstract

The first six postnatal months are a critical period for brain development, marked by rapid changes in functional neural circuits. However, long-term changes in neonatal functional connectome lacks an interpretive imaging indicator for the future development due to the non-linearity characteristics. In this study, we introduce an approach to extract intrinsic brain states from short-term brain dynamics to study the long-term (longitudinal) development. We found a high association (r=0.460) between the co-activated pattern of specific brain state and the acceleration pattern of non-linear development of static functional connectome. The fractional occupancy, self-sustaining probability of this short-term state share the similar age tendency with the long-term change rate within the majority of the function connectome. These findings suggest that short-term brain dynamics could serve as potential biomarkers for predicting the long-term development of functional connectome.

## 1. INTRODUCTION

The first six postnatal months are a critical period for infant development. Functional magnetic resonance imaging (fMRI) studies have shown that macroscale functional neural circuits (e.g. motor, visual, and cognitive networks) undergo rapid and dramatic changes [1, 2]. During this period, infant brain development is characterized by enhanced regional modularity, increased inter-network integration, and growing complexity of the functional connectome. It is gradually more distinguishable for the default mode network, sensorimotor network, and visual network. Multiple factors, including internal genetics and external environmental influences, play substantial roles in the development of the static functional connectome. However, an interpretive imaging indicator to capture the nonlinear development of neonatal brains is lacking, due to many factors involved and their nonlinear interactions in brain development.

Dynamic functional connectivity, a non-static measure of functional activity, describes time-varying functional connectome within a fMRI session. Previous studies have demonstrated its associations with genetics [3], cognition, mental disorders [4] and task-driven characteristics. Neonatal studies have found that established brain states in dynamic connectivity are present by the time of birth and have shown significant differences in transition activity between term-born and preterm infants [5]. Diffusion imaging studies also show the long-term growth in white matter has been found to be associated with the diversity of brain states and the transition ability between them [6]. Dynamic functional connectivity shows its potential as an imaging biomarker for long-term brain development.

In our study, we investigate spontaneous brain dynamics and their role in the longitudinal development of the static connectome. We decompose several transient brain states from spontaneous brain activity using spatial independent component analysis (sICA). By examining the temporal characteristics, we reveal how these transient brain states change with age. Moreover, our findings demonstrate a significant association between specific brain states in the dynamic connectome and the acceleration pattern of change in the static connectome. This suggests that spontaneous brain dynamics within a short term may serve as imaging biomarkers for longer-term change.

## 2. METHOD

### 2.1 Dataset

74 infants (43 males and 31 females) were recruited in prospective longitudinal studies at the Marcus Autism Center in Atlanta, GA, USA. Participants’ parents provided informed written consent, and the Emory University Institutional Review Board approved the research protocol for this study. The infants had a mean gestational age at birth of 39.2 weeks. Infant age at each scan was corrected by gestational age at birth, defined as the age in weeks minus (40 - gestational age in weeks). The subjects used in this study were typically developing infants, with no history of autism in up to 3^rd^ degree relatives and no history of developmental delay in up to 1^st^ degree relatives. Infants with missing gestational age at birth were excluded from further analysis.

Infant scans were acquired at Emory University’s Center for Systems Imaging Core on a 3T Siemens Tim Trio or a 3T Siemens Prisma scanner, using a 32-channel head coil. All infants were scanned during natural sleep. Scans were scheduled for each infant at up to three pseudorandom timepoints between birth and 6 months. More than 1 fMRI run was scanned in a timepoint for an infant. Functional MRI data were acquired using a multiband echo planar imaging (EPI) sequence. The acquisition parameters for the Tim Trio/Prisma scanner were TR = 720/800 ms, TE = 33/37 ms, flip angle = 53°/53°, FOV = 208 × 208 mm, and 2.5/2 mm isotropic spatial resolution.

Data preprocessing followed a previously established functional MRI pipeline [7, 8]. We performed quality controls by selecting infants: 1) with available fMRI scans in at least two longitudinal timepoints 2) head motions<0.65. There are 45 infants with a total of 218 fMRI scans for the longitudinal analysis

### 2.2. Individual intrinsic connectivity networks (ICNs)

We applied the NeuroMark pipeline [9] integrated in the GIFT fMRI toolbox (http://trendscenter.org/software/gift). We used the NeuroMark_fMRI_1.0 template as the spatial priors to extract functional networks for each scan. This template includes 53 replicable ICNs arranged into seven functional domains: subcortical (SCN), auditory (AUD), visual (VIS), sensorimotor (SMN), cognitive-control (CON), default-mode (DMN), and cerebellar (CER) domains. This pipeline is capable of capturing corresponding functional networks while retaining more single-subject variability.

### 2.3. Static and dynamic functional network connectivity

We performed four-steps postprocessing on the network timeseries to remove remaining noise, including: 1) detrending linear, quadratic, and cubic trends, 2) multiple regression of the six realignment parameters and their derivatives, 3) removal and replacement of detected outliers, and 4) band-pass filtering with a cutoff frequency of 0.01 Hz– 0.15 Hz.

We estimated the static functional network connectivity (sFNC) matrix for each scan using the Pearson correlation coefficient between the postprocessed timeseries, yielding a 53 × 53 matrix. The connectivity values were normalized using Fisher-Z transformation.

We next estimated dynamic functional network connectivity (dFNC) from the ICN timeseries. The dFNC was computed via a sliding window approach, which calculated the Pearson correlation coefficient between the timeseries of two ICNs within a 45s window slide in increments of 1 TR. Previous work has shown that 30s to 1 min windows capture reliable brain dynamics [10]. The dynamic connectivity was further normalized into Z score using fisher-z transformation.

### 2.4 Brain state decomposition

The brain states were estimated by implementing the spatial independent component analysis (sICA) on the dFNC estimates. Specifically, we removed the mean functional connectome from dFNC of each scan. Then dFNCs from all participants in all visits with random selected fMRI scans were concatenated along the temporal dimension to form a large matrix. Finally, spatial ICA with ‘infomax’ optimization was applied to this large matrix to obtain *N* brain states patterns. This procedure was performed 20 times, and the most central solution was chosen as the optimal one. The state number *N* here was chosen based on previous infant study[5]. These brain states derived from dFNCs represent the covarying patterns of dynamic brain connectivity. Therefore, each windowed dFNC is characterized by the weighted summarized of the dFNC state patterns.

### 2.5 Brain state temporal features

After the ICA decomposition, we obtain different state patterns and corresponding state timeseries. To evaluate the dynamic properties of the brain states, we introduced two dFNC state features: state fractional occupancy and state transition matrix. State fractional occupancy describes the temporal proportion of each state that dominates the brain. This metric was calculated as the dominant duration of a given state divided by the total scan time. Dominant duration for each state was computed as the proportion of timepoints at which this state had the largest temporal value against the other brain states. The state transition matrix indicates the probability of transitioning from one state to another at the next time point. This metric has the shape of *N × N*, where each row represents the transition probability from one state to different brain states (including itself, the summation of each row is equal to 1). The temporal features correspond to the midpoint of two longitudinal time points are the average of two fMRI scans. To evaluate the tendency of temporal features in the development, we make an age regression for each feature.

### 2.6. Longitudinal change rate of functional connectivity

We first estimated general FNC longitudinal change rate (LCR) using longitudinal data of subjects during the first six month. For one subject, the LCR of one connectivity is the slope of a linear regression between functional connectivity in different visits and subject’s corrected ages. The general LCR is the average of each individual LCR. In addition to the general LCR, we also estimated the instantaneous LCR by the slope of a linear regression between two nearby timepoints divided by time interval. We excluded samples with a time interval greater than three months, as longer intervals may introduce more bias in the estimation of instantaneous LCR. Based on the instantaneous LCR, we are able to investigate whether there is a nonlinear relationship in the development. If the instantaneous LCR is stable during the whole period, there is a linear relationship in the development; otherwise, it does not. The longitudinal change acceleration (LCA) is used for this purpose and estimated as the slope of the linear regression between the instantaneous LCR and corrected age. Their significance was corrected by false discovery rate (FDR).

The LCA reflects the group-level acceleration of the connectivity growth in the brain development. This long-term acceleration will be compared with the pattern of short-term different state patterns, as well as their age tendencies.

## 3. RESULTS

### 3.1. Dynamic functional connectome changes with age in 0-6-month infants

We first decomposed the dFNC brain states from the 0–6-month-old infants. In this work, we used six brain states, the same number in previous study on the infant dynamic pattern [5]. As shown in figure 2, different brain states capture different modularity patterns within the brain. For example, state 1 shows large connectivity between sensorimotor and cognitive control domains, while the state 4 is more focused on visual and cognitive control domains.

**Figure 1.**
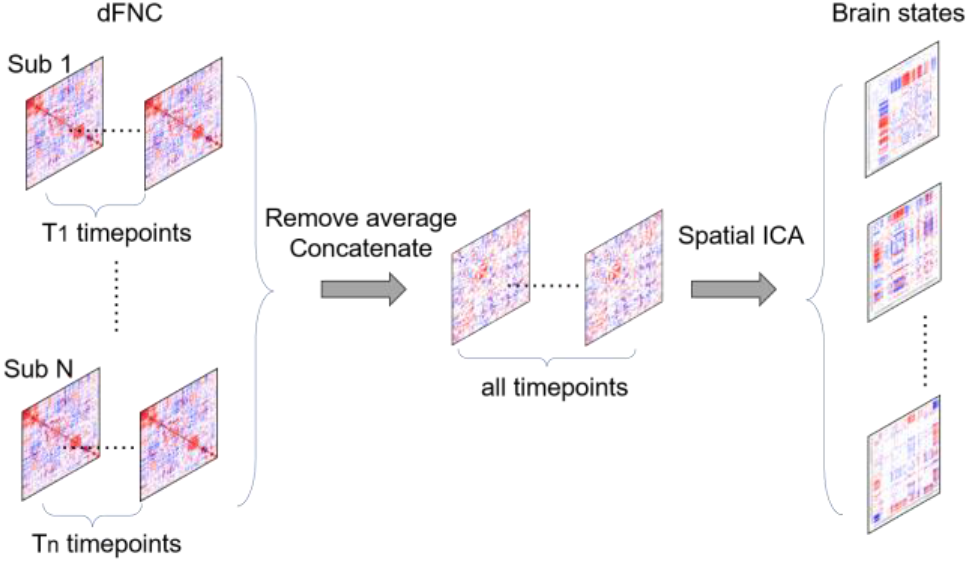
The diagram of brain state decomposition from dFNC using spatial ICA.

**Figure 2.**
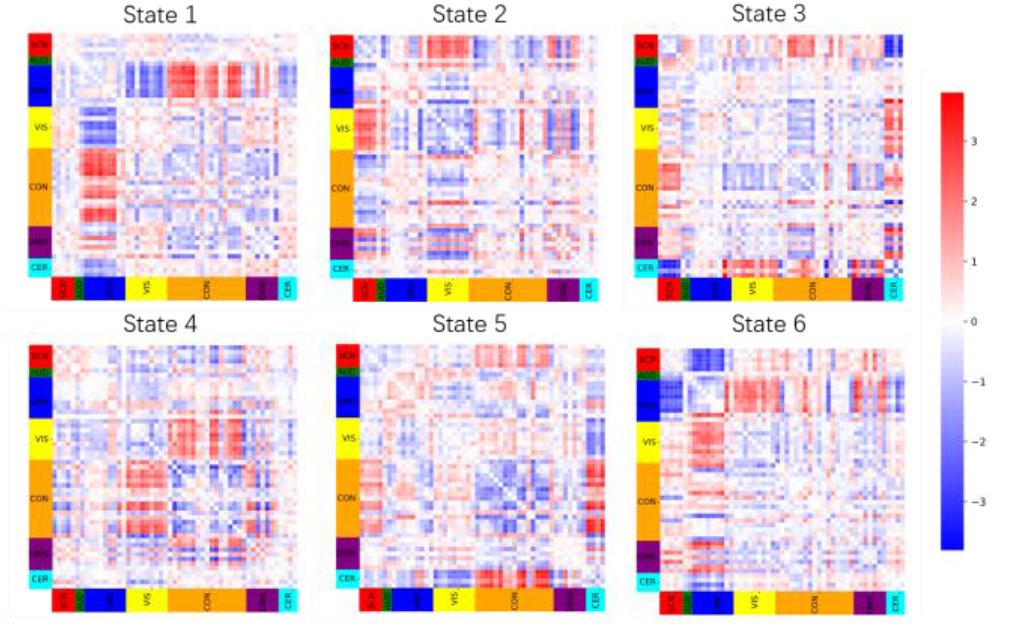
the dFNC brain states across 0-6-month infants.

Moreover, we found that the temporal characteristics of several states significantly correlated with corrected infant age. As shown in Table 1, there is a significant increase in the fractional occupancy of state 4 (r=0.219, p<0.05), and a significant decrease in state 1 (r=-0.176, p<0.05). From the perspective of the state transition matrix (shown in Table 2), state 4 reduced its probability of transitioning to other states as the infants developed, but increased the probability of self-sustaining stays. This aligns with the increasing fractional occupancy of state 4.

**Table 1.** The correlation between corrected age and the fractional occupancy of state time duration for each state. The * indicates the t-test significance with p<0.05, n=219.

| State 1 | State 2 | State 3 | State 4 | State 5 | State 6 |
| --- | --- | --- | --- | --- | --- |
| **-0.176\*** | -0.123 | 0.082 | **0.219\*** | 0.062 | -0.048 |

**Table 2.** The correlation between corrected age and each state transition probability in the state transition matrix. The * indicates the t-test significance with p<0.05, n=219.

|  | S1 | S2 | S3 | S4 | S5 | S6 |
| --- | --- | --- | --- | --- | --- | --- |
| S1 | **-0.211\*** | **0.185\*** | 0.018 | 0.005 | 0.018 | 0.088 |
| S2 | -0.117 | 0.009 | 0.039 | -0.017 | -0.027 | **0.189\*** |
| S3 | 0.012 | -0.033 | 0.031 | 0.047 | -0.011 | -0.075 |
| S4 | **-0.157\*** | -0.005 | **-0.189\*** | **0.165\*** | **-0.17\*** | -0.075 |
| S5 | 0.063 | -0.114 | 0.019 | -0.005 | 0.101 | **-0.189\*** |
| S6 | 0.111 | 0.100 | -0.085 | **0.178\*** | 0.078 | **-0.213\*** |

### 3.2. Nonlinearly brain development of functional connectivity in the first six-month-age infants

We estimated the mean LCR across all 0–6-month-old infants with the assumption that a constant LCR during the first six months of brain development, as shown in Figure 3a. The modularity of the visual network increased dramatically compared to other networks. The regional connectivity between the sensorimotor domain, cognitive control domain, subcortical, and cerebellum domain also increased with corrected age. However, constant hypothesis on LCR was shown to have a significant deviation. The correlation between LCR itself and infant age were significant (p<0.05; FDR corrected) for almost half of connectivity. An example of a connectivity within visual domain, instead of a constant rate of change (red line in Figures 3b and 3d), there is a negative trend (green line in Figure 3d) in the change rate, leading a non-linear growth in connectivity of visual domain (Figure 3b), similar situation for other connectivity within the visual domain (see purple circle in the Figure3c). In contrast, the connectivity between visual and cognitive control domains show acceleration (see green circle in the Figure3c), despite the magnitude of the change itself remains low initially. This may indicate the beginning of another brain remodeling process after gradual maturation of the visual system in infants. Moreover, LCR is also accelerating within the sensorimotor and auditory domains, as well as their interactions with the subcortical domain.

**Figure 3.**
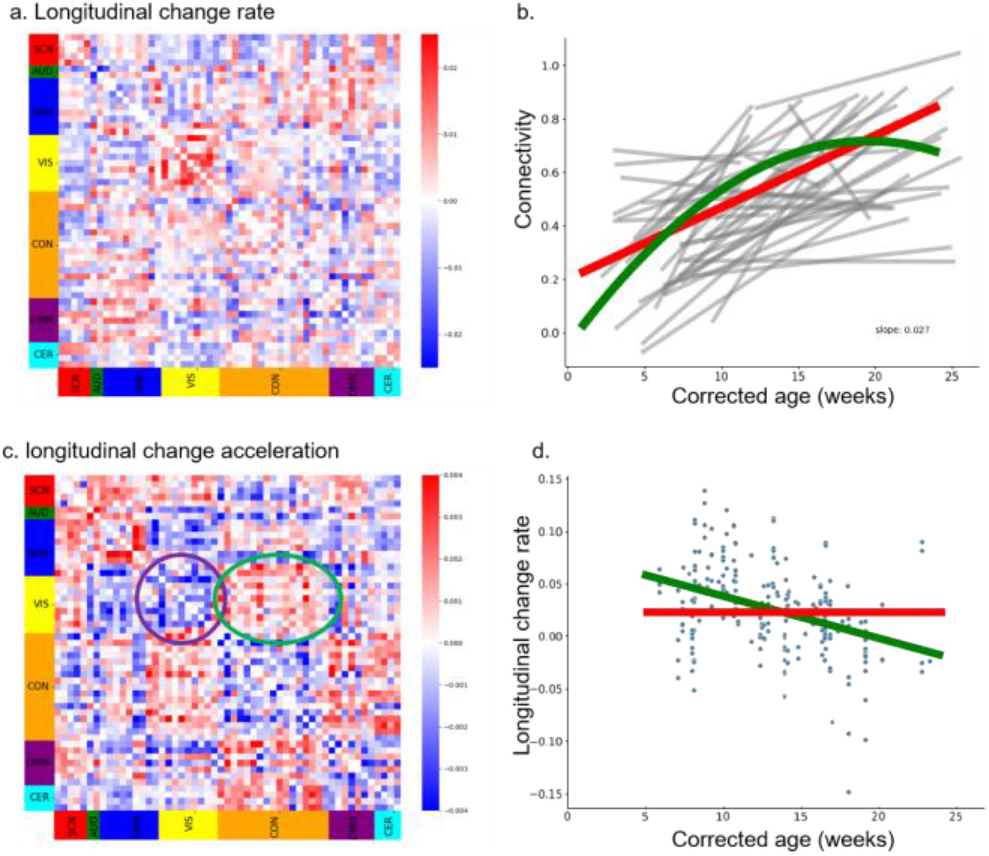
The longitudinal change rate for 0-6-month infants. a: averaged LCR of functional connectome. b: an example of a connectivity within visual domain. c: positive slope for the LCR with corrected age. d: LCR changes with corrected age for the same connectivity with b (p<0.05, 219 paired sample).

### 3.3. Association between the tendency of LCR in the static connectome and the tendency of brain states in the dynamic connectome

Results showed a significant association (r=0.460, p<0.001) between the LCA pattern (shown in Figure 3c) and the modularity pattern of state 4 (shown in Figure 1), and higher than the relationship in the other five state patterns. This correlation is also detected when the total state number is set to 7. It shows a similar age tendency for the temporal feature of specific state, and a significant similarity with the pattern of longitudinal change acceleration. Moreover, the longitudinal change rate in the positive region of state 4 is more positively correlated with the fractional occupancy of the state 4 across subjects. These observations provide evidence that specific characteristics within infant spontaneous brain dynamics are substantial indicators of future longitudinal change rates at the group level.

## 3. DISCUSSION AND CONCLUSION

Our study reveals that the brain dynamics of infants, represented by different brain states, change significantly during the first six months of life. For example, State 1, which reflects strong sensorimotor-cognitive connectivity, decreases in duration over time, while state 4, highlighting visual-cognitive connectivity, increases. This shift indicates that as infants develop, visual-cognitive networks become more dominant, reflecting the rapid maturation of visual functions.

The analysis of the static functional connectome LCR also shows that brain connectivity does not change at a constant rate. Instead, networks like the auditory and motor systems accelerate their development in the first six months, while the change rate of visual network slows down after its early maturation. This suggests that brain development is non-linear, with networks have heterogeneity growth rate.

A significant association was found between the changes in State 4 and the LCR of specific networks, suggesting that specific brain dynamic patterns may help understanding the brain plasticity at younger ages and inform future changes in static brain connectivity.

In summary, our results demonstrate that dynamic brain states and the longitudinal changes in the functional connectome are closely linked during early infancy. The growing dominance of the visual-cognitive brain state, and the varying rates of network development, provide valuable insights into the non-linear progression of brain development.

## 5. ACKNOWLEDGMENTS

The TReNDS/MAC pediatric neuro AI computational core is funded by the Whitehead Foundation

## 6. COMPLIANCE WITH ETHICAL STANDARDS

Participants’ parents provided informed written consent, and the Emory University Institutional Review Board approved the research protocol for this study.

